# Cell-type-specific decoding of Hippo pathway inactivation drives seminiferous epithelial collapse and rete testis hyperplasia in the adult mouse testis

**DOI:** 10.64898/2026.08.05.742520

**Authors:** Laureline Charrier, Laurence Banville, Samuel Gusscott, Nour Abou Nader, Guillaume Saint-Jean, Julie Brind’Amour, Alexandre Boyer

## Abstract

The Hippo pathway is a major regulator of epithelial homeostasis, but its role in the adult testis remains poorly defined. Here, we conditionally deleted *Lats1* and *Lats2* in WT1-expressing cells of the adult mouse testis, targeting both Sertoli cells and rete testis epithelial cells. Deletion of *Lats1/2* caused rapid degeneration of the seminiferous epithelium, leading to near-complete germ cell loss, while simultaneously inducing marked hyperplasia of the rete testis. These tissue alterations were accompanied by vascular remodeling, macrophage accumulation, and extracellular matrix deposition. Cell-type-resolved transcriptomic analyses showed that Sertoli cells and rete testis epithelial cells have distinct baseline identities that shape their response to Hippo pathway inactivation. While both lineages activated shared YAP/TAZ-responsive genes, their broader outputs diverged sharply, with Sertoli cells preferentially activated structural remodeling programs, whereas rete testis epithelial cells robustly activated cell-cycle pathways and lost ciliogenesis-associated features. Regulatory network analysis further identified a conserved Hippo-responsive transcriptional backbone that was redeployed through lineage-specific regulons in each cell type. Together, these findings identify LATS1/2-mediated Hippo signaling as an essential regulator of adult testicular epithelial homeostasis and show that neighboring epithelial lineages decode Hippo pathway disruption through distinct transcriptional programs, resulting in divergent pathological outcomes.

## Introduction

The adult mammalian testis contains highly specialized epithelial compartments that are essential for male fertility. Within the seminiferous tubules, Sertoli cells provide structural, metabolic, and immunological support required for spermatogenesis. They also maintain the blood-testis barrier and contribute to the immune-privileged environment that protects developing germ cells from inflammatory damage (O’Donnell et al., 2022). Following release into the tubular lumen, spermatozoa are transported through the seminiferous tubular fluid to the rete testis, a specialized epithelial conduit that connects seminiferous tubules to the efferent ductules, where much of the luminal fluid is reabsorbed (Clulow et al., 1998; Weigel Munoz et al., 2024). Despite the central role of both the seminiferous tubule epithelium and the rete testis in sperm production and transport, the signaling pathways that maintain these adult epithelial compartments remain incompletely understood.

The Hippo pathway is an evolutionarily conserved signaling cascade that regulates organ size, epithelial architecture, regeneration, and tissue homeostasis (Guo et al., 2025). In its active state, upstream kinases including MST1/2, MAP4Ks, and PKA promote activation of the core kinases LATS1 and LATS2, which phosphorylate and inhibit the transcriptional co-activators YAP and TAZ. When Hippo signaling is inactive, YAP and TAZ accumulate in the nucleus and cooperate with DNA-binding transcription factors, most notably TEAD family proteins (Lopez-Hernandez et al., 2021). Although Hippo signaling has been extensively studied in development and cancer, its role in maintaining differentiated epithelial populations in the adult testis remains poorly defined.

In the developing testis inactivation of *Yap* and *Taz* in Sertoli cells causes modest and transient developmental phenotypes (Levasseur et al., 2017), whereas deletion of *Lats1/2* disrupts testis cord organization (Abou Nader et al., 2024), preventing analysis of their function in mature Sertoli cells. In addition, FSH/PKA signaling can induce YAP phosphorylation in cultured pubertal rat Sertoli cells, suggesting that Hippo signaling remains active after testis development is complete (Sen Sharma et al., 2019). However, whether LATS1/2-dependent Hippo signaling is required in mature Sertoli cell maintenance has not been tested, while the role of Hippo signaling in rete testis epithelial cells remains unexplored.

Here, we used a tamoxifen-inducible *Wt1*^CreERT2^ model to delete *Lats1* and *Lats2* in WT1-expressing cells of the adult mouse testis. We show that Hippo pathway inactivation causes rapid seminiferous epithelial degeneration together with marked rete testis hyperplasia. By combining whole-testis and cell-type-resolved transcriptomic analyses, we further show that Sertoli and rete testis epithelial cells respond to *Lats1/2* deletion through a shared YAP/TAZ-associated transcriptional core that is redirected into cell type-specific remodeling programs. These findings show that Hippo pathway inactivation elicits divergent pathological responses in neighboring adult testicular epithelial lineages and highlight cell identity as a major determinant of this response.

## Results

### *Lats1/2* deletion in WT1-expressing adult testicular epithelial cells causes seminiferous tubule degeneration and rete testis hyperplasia

To investigate the role of Hippo signaling in adult testicular homeostasis, we generated *Lats1*^fl/fl^; *Lats2*^fl/fl^; *Wt1*^CreERT2/+^ mice (hereafter *L1L2*-cKO), and induced recombination in Sertoli and rete testis epithelial cells at 8 weeks of age by tamoxifen administration for 5 consecutive days (**Figure 1A**). Control mice consisted of tamoxifen-treated *Lats1*^fl/fl^; *Lats2*^fl/fl^ littermates (hereafter *L1L2*-Ctl). Only tamoxifen-treated mutant mice developed testicular abnormalities. By day 12 after tamoxifen injection (D12), *L1L2*-cKO mice displayed systemic morbidity and renal dysfunction, and this time point was therefore selected as the experimental endpoint.

**Figure 1.**
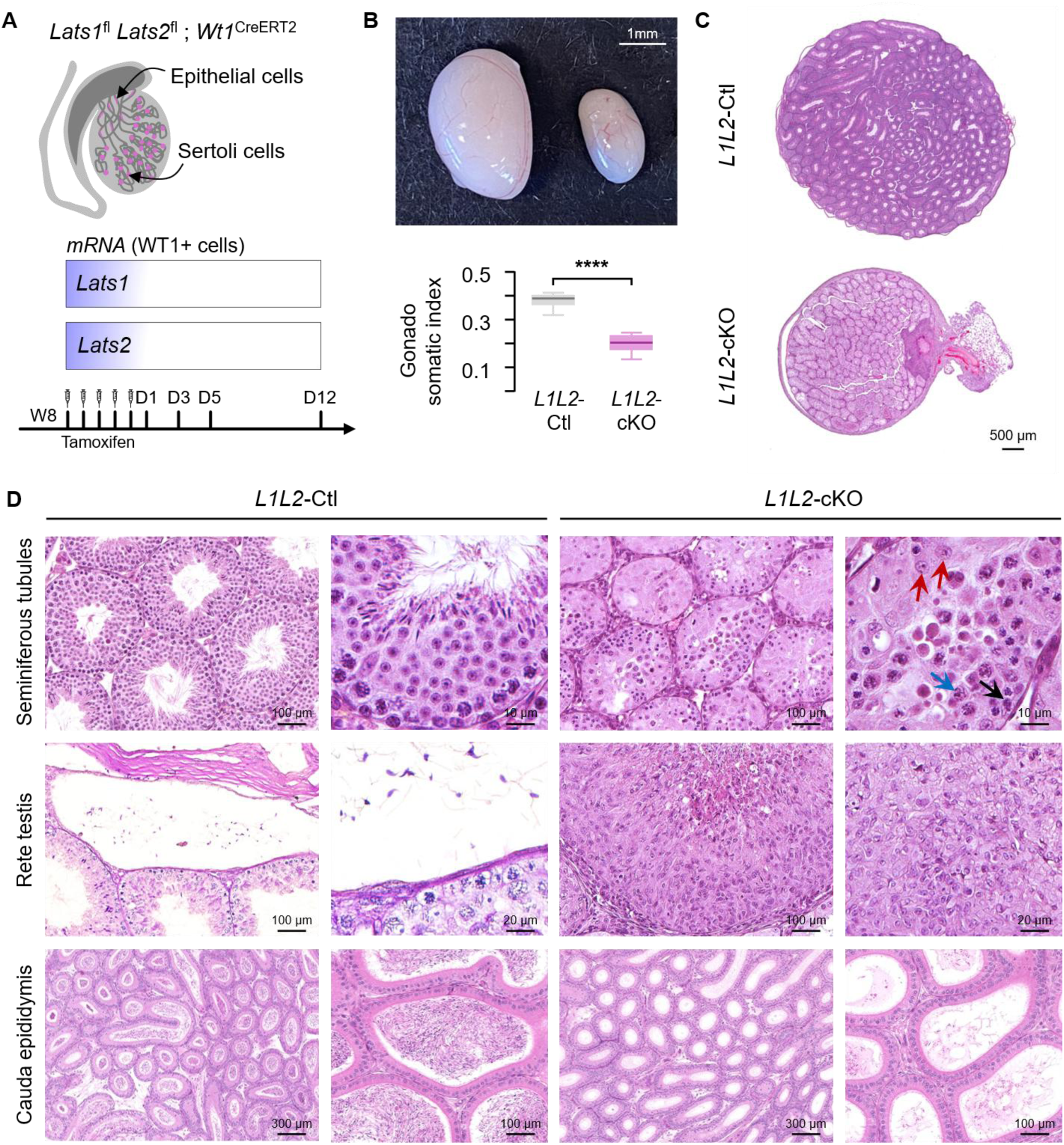
Tamoxifen-induced deletion of *Lats1* and *Lats2* in Sertoli cells leads to severe testicular disorganization. **(A)** Experimental design for tamoxifen-induced deletion of *Lats1* and *Lats2* using *Wt1*^CreERT2^ driver. **(B)** Representative macroscopic images and gonadosomatic index of *L1L2*-Ctl and *L1L2*-cKO testes at D12 (n = 6 per group, ****p < 0.0001, Student’s t test). **(C)** H&E staining of *L1L2*-Ctl and *L1L2*-cKO testes at D12. Scale bars, 500 µm. **(D)** Higher-magnification images of seminiferous tubules, rete testis and cauda epididymis from *L1L2*-Ctl and *L1L2*-cKO testes at D12. Arrows indicate spermatogonia (black), spermatocytes (blue), and Sertoli cells (red). Scale bars, as indicated.

At D12, *L1L2-*cKO testes were markedly reduced in size and exhibited a twofold decrease in gonadosomatic index relative to controls (**Figure 1B**). Histopathological analysis revealed severe disruption of seminiferous tubule architecture, characterized by altered Sertoli cell morphology, including loss of polarity and cytoplasmic expansion, accompanied by intraluminal cellular debris presence, and extensive germ cell loss. In most tubules, only spermatogonia and early primary spermatocyte remained (**Figure 1C, 1D**). In parallel, the rete testis showed pronounced epithelial dysplasia, with epithelial multilayering, luminal narrowing and occlusion, and accumulation of intraluminal debris (**Figure 1C, 1D**). Consistent with profound impairment of spermatogenesis and/or sperm transport, the cauda epididymis of *L1L2*-cKO animals lacked spermatozoa (**Figure 1D**). No comparable abnormalities were observed in *L1L2*-Ctl testes (**Figure 1D**).

To confirm Hippo pathway inactivation, we assessed YAP/TAZ phosphorylation in control and mutant testes. In *L1L2*-Ctl testes, YAP, TAZ and their phosphorylated forms (pYAP/pTAZ) were detected in both Sertoli and rete testis epithelial cells (**Figure S1A**). In contrast, *L1L2-*cKO testes retained YAP/TAZ protein expression but showed loss of pYAP and pTAZ staining in the targeted cells, confirming effective inactivation of LATS1/2 signaling (**Figure S1A**).

We next examined the temporal progression of the phenotype. In contrast to D12, an increase in gonadosomatic index was evident by D3 and persisted at D5 (**Figure S1B, S1C**). Subtle thinning of the seminiferous epithelium was detectable at D1, despite preserved overall testicular architecture, whereas no morphological changes were observed in the rete testis at this stage (**Figure S1D, S1E**). By D3, seminiferous tubule epithelium atrophy and loss of elongated spermatids were evident, and epithelial hyperplasia became detectable in the rete testis. By D5, germ cell depletion had progressed to spermatocytes, and rete testis was occluded (**Figure S1D, S1E**). Because defects in the seminiferous epithelium precede complete rete testis obstruction, these results support the idea that LATS1/2 contributes directly to seminiferous tubule homeostasis, while not excluding additional secondary effects resulting from altered rete testis function.

### *Lats1/2* deletion disrupts spermatogenesis and remodeling of the adult testicular microenvironment

To define the molecular consequences of *Lats1/2* deletion, we performed RNA sequencing on whole testes from *L1L2*-Ctl and *L1L2*-cKO mice. Differential expression analysis revealed widespread transcriptional remodeling in mutant testes, with 1,717 upregulated and 1,354 downregulated genes (**Figure 2A**). Downregulated genes were associated with germ cell differentiation, including male gonad development, single fertilization and sperm motility, indicating impairment of spermatogenesis (**Figure 2B**). Corroborating the histological phenotype, analysis of defined stage-specific germ cell signatures further showed relative preservation of spermatogonial and early spermatocytes markers together with progressive loss of transcripts associated with pachytene spermatocytes, round spermatids, and elongated spermatids (**Figure 2C**).

**Figure 2.**
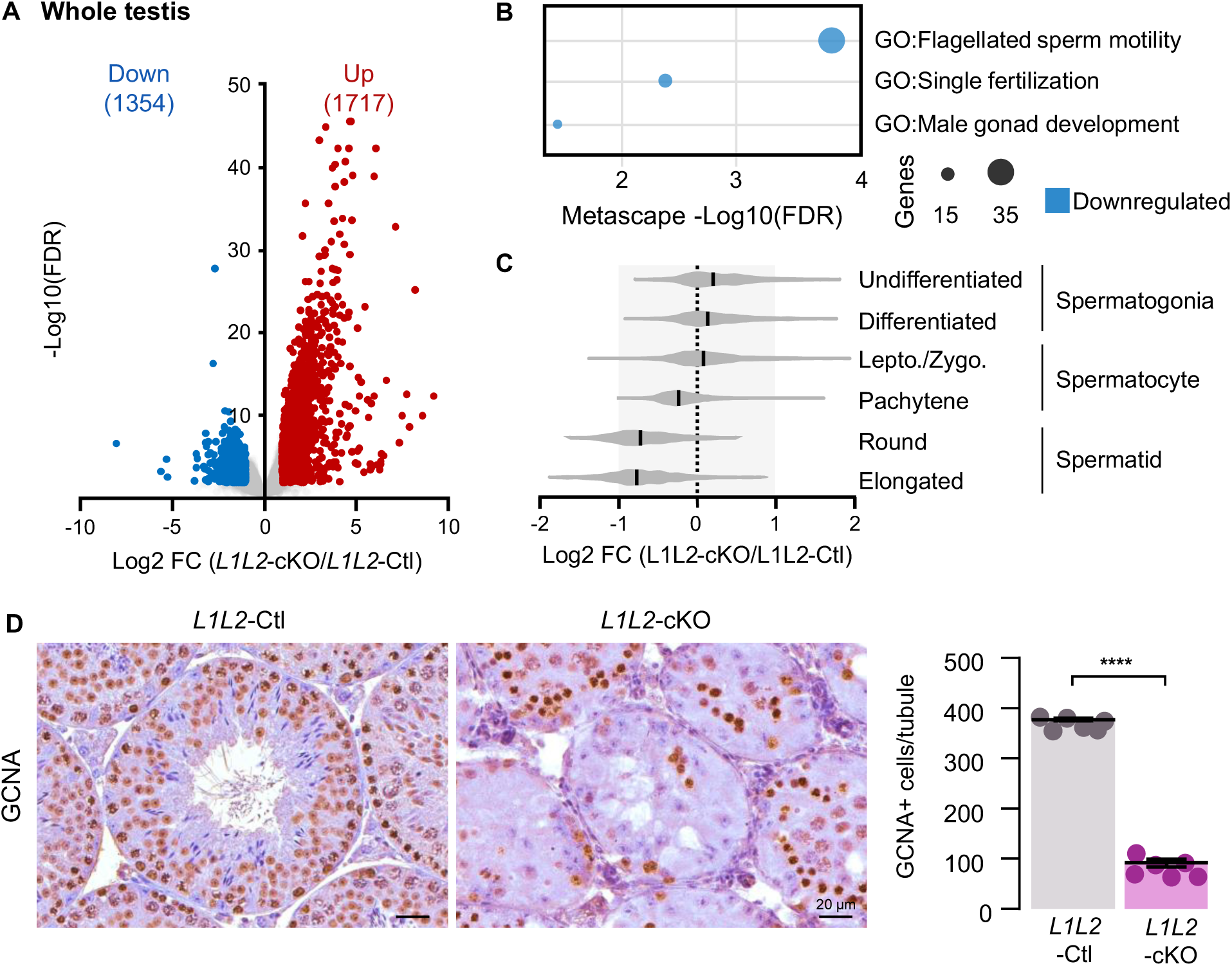
*Lats1/2* deletion disrupts spermatogenesis and germ cell organization. **(A)** Volcano plot showing differentially expressed genes between *L1L2*-cKO and *L1L2*-Ctl in whole-testis RNA-seq. **(B)** Gene Ontology (GO) enrichment of downregulated genes in *L1L2*-cKO testes. **(C)** Violin plots of germ cell subtype medians, based on theoretical versus observed distributions across six spermatogenic stages (Jung et al., 2019). Two-tailed Wilcoxon Signed Rank tests (****p < 0.0001 for all stages). Data represent median ± 95% confidence interval. **(D)** IHC for GCNA in *L1L2*-Ctl and *L1L2*-cKO testes. Scale bars, 20 µm. Quantification of GCNA⁺ cells per seminiferous tubule (n = 6; ****p < 0.0001, Student’s t test).

Consistent with these transcriptomic changes, immunostaining for the germ cell marker GCNA revealed an approximately four-fold reduction in germ cell number per seminiferous tubule in *L1L2*-cKO testes (**Figure 2D**). In control testes, germ cells formed multiple orderly layers along the seminiferous epithelium. In contrast, GCNA-positive cells in mutant tubules were sparse, disorganized, and frequently displaced toward the lumen, with relatively few cells remaining at the basement membrane (**Figure 2D**).

Because germ cell organization depends on Sertoli cell structure and junctional integrity, we next examined markers of seminiferous epithelial architecture. At D5, Connexin43, beta-catenin and vimentin showed largely comparable localization in control and mutant testes, suggesting that major junctional disruption had not yet occurred (**Figure S2**). By D12, however, Connexin43, beta-catenin and vimentin staining were altered, consistent with progressive Sertoli cell remodeling and cytoskeletal disorganization (**Figure S2**). These findings suggest that loss of germ cell organization is initiated before overt collapse of Sertoli cell junctional architecture.

Whole-testis transcriptomic analysis also revealed marked induction of pathways associated with vascular remodeling, inflammatory responses, wound response, and cytokine production (**Figure 3A**). Consistent with this observation, mutant testes displayed increased PECAM1-positive vascular structures and prominent accumulation of CD204-positive macrophages around degenerating seminiferous tubules and within the expanded rete testis epithelium (**Figure 3B**). These responses were already detectable at D5 (**Figure S3A**). In addition, Picrosirius Red staining revealed increased collagen deposition around seminiferous tubules and within the hyperplastic rete testis epithelium, consistent with extracellular matrix remodeling and fibrosis often associated with chronic inflammatory/immune response (**Figure S3B**).

**Figure 3.**
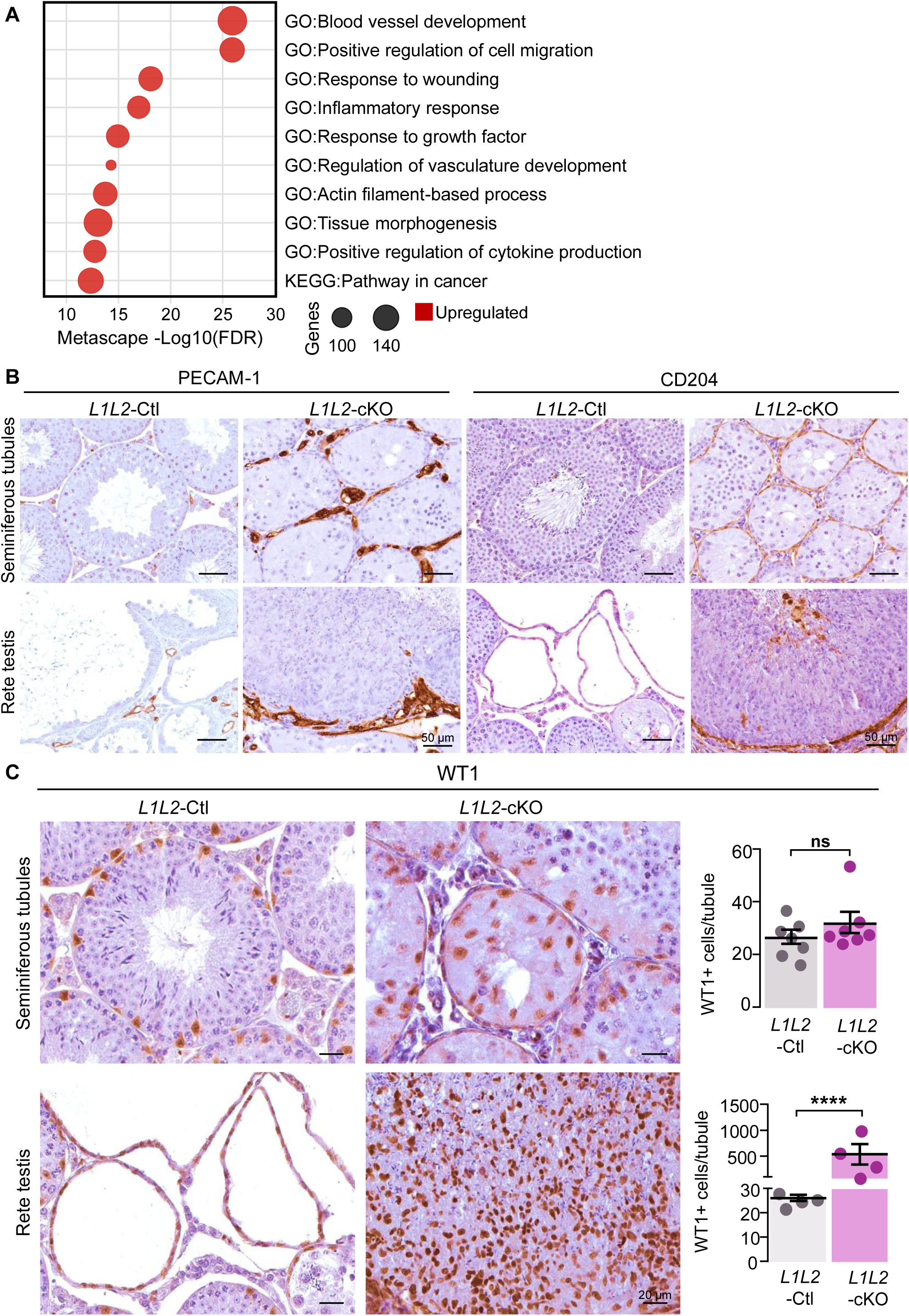
*Lats1/2* deletion remodels vascular and immune microenvironments. **(A)** GO enrichment of upregulated genes in *L1L2*-cKO testes. **(B)** IHC for PECAM 1 (endothelial cells) and CD204 (macrophages) in *L1L2*-Ctl and *L1L2-*cKO testes at D12. Scale bars, 50µm. **(C)** IHC for WT1 in seminiferous tubules (top) and rete testis (bottom) in *L1L2*-Ctl and *L1L2*-cKO testes at D12. Quantification of WT1+ cells per seminiferous tubule (n = 7 per group; **** p < 0.0001, Student’s t test). Scale bars, 20 µm.

Despite being exposed to the same upstream Hippo perturbation, Sertoli cells and rete testis epithelial cells exhibited distinct cellular responses. A second group of enriched pathways was associated with tissue remodeling and proliferative growth, consistent with extensive structural reorganization of the rete testis epithelium (**Figure 3A**). PCNA staining revealed robust proliferative activity in the rete testis epithelial cells of *L1L2*-cKO testes, whereas Sertoli cells remained PCNA-negative, with only PCNA-positive germ cells detected within the seminiferous tubules (**Figure S3C**). Quantification of WT1-positive cells showed an approximately 20-fold increase in the rete testis of mutant testes, whereas WT1-positive cell numbers within seminiferous tubules were not significantly changed (**Figure 3C**). Moreover, rete testis epithelial cells in *L1L2*-cKO testes showed increased vimentin expression while beta-catenin remained membranous, with staining patterns consistent with acquisition of mesenchymal-associated features while retaining epithelial characteristics (**Figure S3C**). Together, these data indicate that *Lats1/2* deletion drives coordinated remodeling of the adult testis, coupling seminiferous epithelial degeneration to vascular, immune, and extracellular responses, while eliciting sharply divergent outcomes in neighboring WT1-positive epithelial populations.

### Sertoli cells and rete testis epithelial cells display distinct baseline identities and activate both shared and specific transcriptional responses after *Lats1/2* deletion

To define cell-type-specific responses to Hippo pathway inactivation, we generated *Rosa*^mTmG/mTmG^; *Lats1*^fl/fl^; *Lats2*^fl/fl^; *Wt1*^CreERT2/+^ mutant mice and *Rosa*^mTmG/mTmG^; *Wt1*^CreERT2/+^ control mice in which tamoxifen-induced Cre activity labeled WT1-expressing cells with eGFP for lineage tracing and cell isolation. Following microdissection, tissue dissociation, and fluorescence-activated cell sorting at D12 after tamoxifen treatment, bulk RNA sequencing was performed on four populations: Sertoli control (S-Ctl), Sertoli mutant (S-cKO), rete testis epithelial control (E-Ctl) and rete testis epithelial mutant (E-cKO) (**Figure 4A, S4A**). Principal component analysis of normalized expression profiles confirmed separation by both cell type and genotype (**Figure S4B**), and comparison of control cell populations revealed marked baseline differences between the two WT1-positive epithelial lineages, with 962 genes enriched in rete testis epithelial cells and 654 enriched in Sertoli cells (**Figure S4C**). Analysis of lineage marker expression supported enrichment of the sorted control populations (**Figure S4D**). Thus, even before perturbation, these neighboring epithelial compartments are defined by distinct transcriptional identities. Efficient excision of *Lats1* and *Lats2* exons was confirmed in both mutant cell populations (**Figure S4E**).

**Figure 4.**
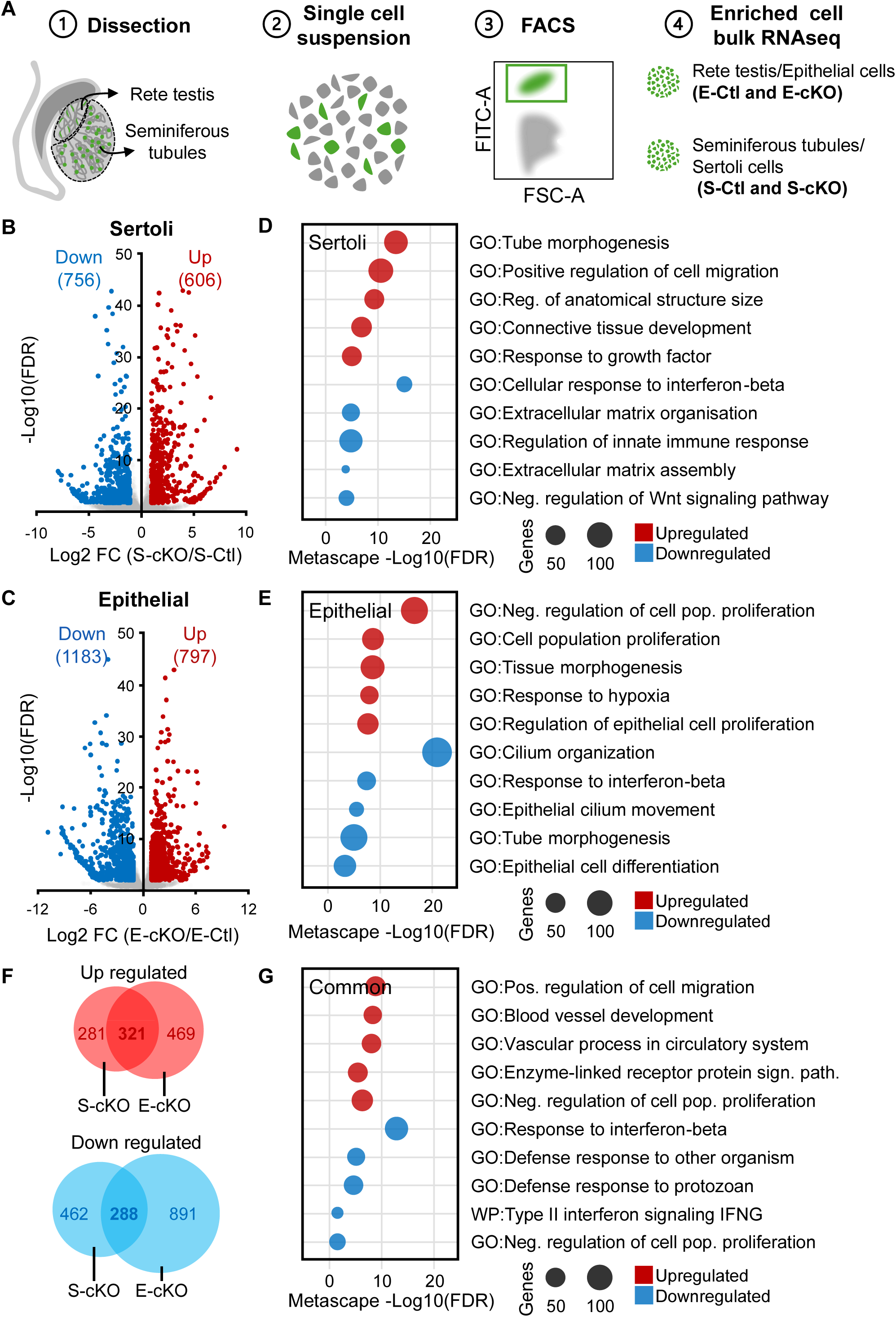
*Lats1/2* deletion triggers distinct transcriptional responses in Sertoli and rete testis epithelial cells. **(A)** Experimental design for enrichment of WT1+ Sertoli and rete testis epithelial cells for RNA-seq. **(B)** Volcano plot showing DEGs in Sertoli cells (S-cKO *vs* S-Ctl). **(C)** Volcano plot of DEGs in rete testis epithelial cells (E-cKO *vs* E-Ctl). **(D)** GO enrichment of DEGs in Sertoli cells (S-cKO *vs* S-Ctl). **(E)** GO enrichment of DEGs in rete testis epithelial cells (E-cKO *vs* E-Ctl). **(F)** Overlap of upregulated and downregulated genes between the two lineages. **(G)** GO enrichment analysis of shared transcriptional responses in Sertoli and rete testis epithelial cells.

Deletion of *Lats1/2* induced extensive but divergent transcriptional remodeling in each lineage (**Figure 4B-4C**). In Sertoli cells, upregulated genes were enriched for morphogenesis, migration, and connective tissue development consistent with structural remodeling, whereas downregulated genes reflected reduced homeostatic and signaling functions, including pathways linked to Wnt signaling and extracellular matrix organization (**Figure 4D**). In rete testis epithelial cells, upregulated genes were strongly enriched for cell-cycle progression, proliferation, hypoxia-related responses, and epithelial migration, while downregulated genes suggested loss of differentiated epithelial features, most notably ciliogenesis (**Figure 4E**).

Although the two WT1-positive testicular lineages respond differently, overlap analysis identified 321 commonly upregulated genes and 288 commonly downregulated genes (**Figure 4F**). Shared upregulated genes were enriched for cell migration, vascular development, and tissue remodeling, whereas shared downregulated genes were enriched for immune-related and interferon-associated pathways, as well as negative regulators of proliferation (**Figure 4G**).

We next examined a curated set of know YAP/TAZ-responsive genes. This analysis identified 35 genes commonly altered in both cell types, alongside 16 Sertoli-specific and 42 rete testis epithelial-specific subsets (**Figure S4F**). Canonical YAP/TAZ targets, including *Ankrd1*, *Ctgf*, and *Cyr61*, were induced in both lineages, supporting activation of a common Hippo-responsive transcriptional core (**Figure S4G**). Thus, Hippo pathway inactivation engages shared YAP/TAZ-associated outputs in Sertoli cells and rete testis epithelial cells, but these shared outputs are redirected into distinct lineage-specific programs.

### A shared Hippo-responsive regulatory core is rewired into distinct transcriptional modules

To investigate how *Lats1/2* deletion produces divergent transcriptional outcomes in Sertoli and rete testis epithelial cells, we inferred transcription factor activity using VIPER and DoRothEA regulons. Comparison of mutant and control populations identified 25 transcription factors with concordant activity changes in both lineages including YAP/TAZ binding partners TEAD1, TEAD4 and HIF1A, defining a shared *Lats1/2*-responsive regulatory core (**Figure 5A**). In parallel, 33 transcription factors displayed Sertoli-specific activity changes and 21 displayed rete testis epithelial-specific activity changes, indicating substantial lineage-specific regulatory remodeling (**Figure 5A**).

**Figure 5.**
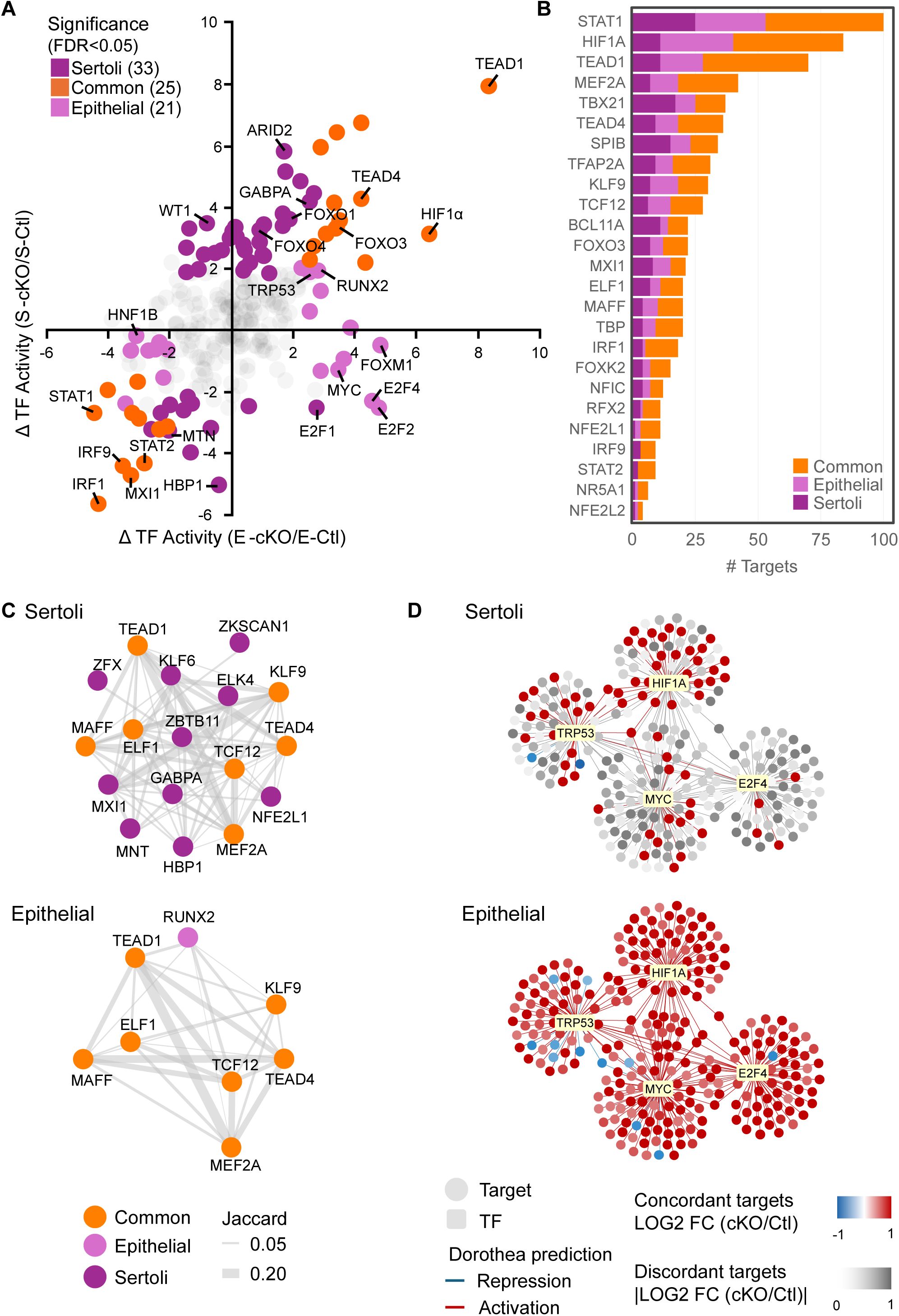
A core YAP/TAZ-responsive program drives shared and specific responses to Hippo pathway inactivation. **(A)** Scatter plot illustrating differential VIPER-inferred TF activity (cKO/Ctl) in Sertoli and rete testis epithelial cells. TFs are colored by specificity: Sertoli (purple), epithelial (pink) or shared (orange). **(B)** Similarity and regulon size of the 25 TFS displaying concordant activity changes in both cell types, colored by target specificity: Sertoli (purple), epithelial (pink), or shared (orange). **(C)** Regulatory networks of Hippo pathways-associated factors TFS with overlapping regulons, reconstructed independently for Sertoli and rete testis epithelial cells. Edge weights correspond to regulon overlap (Jaccard index). TFs colored as (A). **(D)** Regulatory networks centered on proliferative module. Node shape indicates TFs (rectangles) and targets (circles); edge color represents the predicted mode of regulation (DoRothEA). Target nodes are colored by concordance with differential gene expression (LOG2FC), with discordant targets in gray.

Furthermore, concordant changes in inferred transcription factor activity did not imply identical downstream outputs. Among transcription factors showing similar shifts in activity for both cell type, some were associated with overlapping target gene sets, whereas others were linked to largely distinct regulons depending on cellular context (**Figure 5B**). This suggests that common Hippo-responsive regulators are interpreted through cell-type-specific regulatory architectures rather than through a uniform transcriptional program.

Module-based analysis further highlighted distinct organization of the regulatory response in each cell type (**Figure S5A-S5F**). In Sertoli cells, regulon-based modules were enriched with structural remodeling, apoptosis, metabolism, antiviral defense, and cytoskeletal organization (**Figure S5C**). In rete testis epithelial cells, modules were linked to cell-cycle progression, GTPase signaling, adhesion, lipid metabolism, and immune-related processes (**Figure S5F**). Within the Hippo-responsive module, both cell types shared inferred activity of YAP/TAZ-associated factors including TEAD1, TEAD4, MAFF, ELF1, TCF12, MEF2A, and KLF9 (**Figure 5C**). However, each cell type also engaged distinct additional regulators, Sertoli cells recruited a broader set of cell type-specific factors, whereas rete testis epithelial cells recruited a more limited set of factors, including the selective engagement of RUNX2 (**Figure 5C**).

To better understand the proliferative phenotype of the rete testis epithelium, we examined the cell type-specific proliferative module identified in mutant epithelial cells. Among the transcription factors with differential lineage behavior, MYC and E2F4 displayed increased inferred activity in rete testis epithelial cells and reduced activity in Sertoli cells. TRP53 and HIF1A also emerged as prominent regulators of

large rete testis epithelial cell regulons. These factors converged within a shared proliferation-associated module, providing a plausible regulatory basis for the hyperplastic response of the rete testis epithelium following *Lats1/2* deletion (**Figure 5D**). Together, these analyses support a model in which Hippo pathway inactivation activates a conserved YAP/TAZ-associated regulatory backbone that is subsequently rewired into distinct lineage-specific transcriptional programs.

## Discussion

To this day, the role of Hippo signaling in the adult testis has remained largely unexplored. In this study, using inducible deletion of *Lats1* and *Lats2* in WT1-expressing cells, we show that LATS1/2-dependent Hippo signaling is required to preserve epithelial homeostasis in seminiferous tubules and rete testis of the adult mouse testis. Hippo pathway inactivation triggered two striking and divergent outcomes in neighboring epithelial compartments: seminiferous epithelial degeneration with germ cell loss, and proliferative expansion of the rete testis epithelium. Cell-type-resolved transcriptomic analyses further showed that these distinct phenotypes arise from a combination of shared YAP/TAZ-associated responses and lineage-specific regulatory programs.

A central finding of this study is that the same upstream perturbation does not produce a uniform epithelial response. Although Sertoli cells and rete testis epithelial cells both activated canonical YAP/TAZ-responsive genes, their broader transcriptional outputs diverged sharply. While both cell types induced programs linked to microenvironment modifications, including angiogenesis and macrophage recruitment, Sertoli cells preferentially induced programs linked to extracellular matrix organization, structural remodeling, and tissue stress, whereas rete testis epithelial cells robustly activated cell-cycle programs and lost features of differentiated epithelial identity, including ciliogenesis-associated gene expression. This is consistent with the broader view that YAP/TAZ activity does not dictate a single stereotyped biological outcome but instead acts through a context-dependent transcriptional framework shaped by cell identity, chromatin state, and pre-existing regulatory architecture (Moya & Halder, 2019; Panciera et al., 2017). Our analyses of inferred transcription factor activity support this model by showing that even when both lineages engage a partially shared Hippo-responsive regulatory core, the downstream regulon structure remains highly lineage restricted.

The response of adult Sertoli cells also highlights an important distinction between developmental and adult Hippo pathway function. *Lats1/2* deletion in immature Sertoli cells was previously shown to cause disorganization and transient overgrowth of the testis cords (Abou Nader et al., 2024), no comparable transient increase in proliferation was observed in mature Sertoli cells after Lats1/2 deletion. Instead, they remained post-mitotic and underwent progressive structural deterioration, associated with altered junctional organization, extracellular matrix remodeling, and collapse of spermatogenic support, suggesting that Sertoli cell maturation could alter how Hippo pathway disruption is decoded.

The inflammatory and stromal remodeling phenotype observed following *Lats1/2* deletion is consistent with other previously reported consequences of Hippo pathway inactivation in multiple tissues (Abou Nader et al., 2024; Abou Nader et al., 2025; Mooring et al., 2020; Turinsky et al., 2025) and further underscores the essential role of Hippo signaling in maintaining the adult seminiferous microenvironment. Sertoli cells are key regulators of the immune-privileged environment of the testis, and disruption of this niche is well recognized to impair spermatogenesis (Hedger, 2011; Zhao et al., 2014). Accordingly, vascular expansion, macrophage accumulation, collagen deposition, and activation of inflammatory and wound-response pathways observed following *Lats1/2* deletion are likely to amplify tissue dysfunction and contribute to germ cell depletion. Indeed, macrophage-derived cytokines have been shown to disrupt Sertoli cell junctions and compromise blood-testis barrier, whereas extracellular matrix remodeling interferes with the dynamic junctional restructuring events required for germ cell movements during spermatogenesis (Siu & Cheng, 2008; Theas, 2018). Thus, the progressive disruption of junction-associated markers observed in Sertoli cells likely reflects intrinsic consequences of *Lats1/2* deletion and secondary alteration of the surrounding microenvironment.

In contrast, rete testis epithelial cells exhibited a more classic proliferative response to Hippo pathway inactivation. These cells proliferate rapidly with strong activation of cell-cycle-associated transcriptional programs leading to hyperplasia of the rete testis compartment. This behavior is broadly consistent with the well-established growth-restrictive function of LATS1/2 in adult epithelia and with studies showing that Hippo pathway disruption can promote proliferative expansion and epithelial plasticity in other organs (Carter et al., 2021; Totaro et al., 2018; Zhu et al., 2025). Although the rete testis epithelium displayed multilayering, increased proliferation, and altered expression of epithelial and mesenchymal-associated markers, the present data do not demonstrate malignant transformation.

One notable feature of the rete testis response was the coordinated repression of ciliogenesis-associated genes. Rather than representing isolated transcriptional changes, these alterations point to collapse of a broader ciliogenic differentiation program. This is potentially important because primary cilia are increasingly recognized as regulators of epithelial quiescence, differentiation, and cell-cycle control, and ciliary disassembly is often linked to proliferative entry (Basten & Giles, 2013; Goetz & Anderson, 2010; Wheway et al., 2018). Together, the loss of ciliogenic identity, the reciprocal gain of vimentin expression and sustained proliferative activity suggest that *Lats1/2* deletion drives partial dedifferentiation of rete testis epithelial cells toward a less specialized, proliferative state, rather than transdifferentiation into an alternative differentiated lineage. Whether loss of ciliogenic identity is a direct consequence of Hippo pathway disruption or a secondary adaptation to proliferative activation remains unresolved, but this feature further supports the conclusion that *Lats1/2* deletion drives a marked shift away from normal rete testis epithelial differentiation.

Our analysis of transcription factor activity provides a mechanistic framework for understanding these divergent outcomes. Shared activation of TEAD-associated and other Hippo-responsive factors was accompanied by major differences in regulon composition and by selective mobilization of proliferative and stress-response regulators in rete testis epithelial cells. Among these, MYC, E2F4, HIF1A and TRP53 emerged as highly connected regulators within the inferred rete testis network, with MYC- and E2F-associated modules have been repeatedly implicated in YAP-driven proliferative outputs in other systems (Cordenonsi et al., 2011; Misra & Irvine, 2018; Totaro et al., 2018; Zanconato et al., 2015), and their selective engagement in the rete testis provides a plausible regulatory basis for the hyperplastic phenotype observed here. Other cell-cycle-associated regulators including FOXM1 and E2F2, also displayed increased activity, further supporting the activation of a proliferative transcriptional program in rete testis epithelial cells. The emergence of HIF1A and TRP53 within this proliferative module is consistent with the metabolic and replicative stress associated with rapid epithelial expansion: HIF1A likely reflects hypoxia-driven adaptation and compensatory angiogenesis typical of rapidly expanding tissues (Wicks & Semenza, 2022), while TRP53 activity, despite its canonical tumor-suppressive role, more plausibly reflects a checkpoint response to replication stress than a shift toward a tumor-suppressive state (Engeland, 2022), consistent with the absence of malignant features in this model. At the same time, these inferences should be interpreted cautiously, as they are based on transcriptome-derived regulatory activity rather than direct measurement of chromatin occupancy or factor binding. Even so, the data strongly support a model in which Hippo pathway inactivation activates a conserved YAP/TAZ-associated regulatory backbone that is subsequently interpreted through cell type-specific transcriptional architectures.

Why Sertoli cells and rete testis epithelial cells should differ so markedly remains an open question. One plausible contributing factor is that Sertoli cells are terminally post-mitotic and depend on a stable junctional architecture to sustain the blood-testis barrier, a function likely requiring durable epigenetic repression of cell-cycle programs that loss of Hippo signaling alone may not be sufficient to overcome (Ahmed, 2026; Walker, 2003). In contrast, rete testis epithelial cells may retain a proliferative competence, allowing Hippo pathway disruption to more readily engage cell-cycle programs (Major et al., 2021). This distinction remains speculative and will require direct comparison of chromatin accessibility and cell-cycle regulatory architecture between these two cell types.

Several limitations of this study should be considered. First, the *Wt1*^CreERT2^ model targets both Sertoli cells and rete testis epithelial cells, preventing strict separation of primary and secondary effects across compartments. For example, rete testis obstruction could exacerbate seminiferous damage. However, the early appearance of seminiferous epithelial defects before complete luminal occlusion supports a direct requirement for Hippo signaling in seminiferous tubule homeostasis. Second, the regulatory inferences presented here are based on transcriptomic analyses and inferred transcription factor activity rather than direct perturbation or chromatin-based assays. Third, the relatively short experimental window imposed by extra-testicular morbidity limits assessment of longer-term adaptation or progression of the lesions. Future studies using compartment-restricted models and direct tests of YAP/TAZ-TEAD dependency will be important to refine these mechanisms.

Overall, our findings establish LATS1/2-mediated Hippo signaling as an essential regulator of adult testicular epithelial homeostasis. More broadly, they position the adult testis as a useful system for understanding how neighboring epithelial lineages decode the same signaling perturbation through distinct regulatory states. In this context, the principal conceptual advance of the study is not only that Hippo signaling is required in the adult testis, but that its disruption reveals how lineage identity channels a conserved YAP/TAZ-responsive program toward divergent pathological outcomes: structural collapse in Sertoli cells and hyperplastic epithelial remodeling in the rete testis.

## Resource availability

### Lead contact

Further information and requests for resources and reagents should be directed to and will be fulfilled by the lead contact, Alexandre Boyer

## Materials availability

This study did not generate new unique reagents.

## Data and code availability

RNA-sequencing data generated in this study have been deposited in the Gene Expression Omnibus (GEO) under accession number GSE316188. All other data supporting the findings of this study are available from the corresponding author.

## Acknowledgements

The authors would like to thank Dr Randy L. Johnson (M.D. Anderson Cancer Center, Houston Tx) for generously providing the Lats1/2 floxed mice and Mélanie Lehoux for FACS sorting. This work was supported by a Discovery Grant (RGPIN-2020-05230) and a Discovery Accelerator Supplement (RGPAS-2020-00026) from the Natural Sciences and Engineering Research Council of Canada (NSERC) to A.B. as well as a Discovery Grant (RGPIN-2023-03824) from NSERC and a Project Grant (202303PJT) from the Canadian Institute for Health Research (CIHR) to J.B.

## Authors contributions

Methodology, Formal Analysis, Investigation, Original Draft, Writing, Review & Editing, L.C.; Investigation, N. A. N.; Investigation, L.B.; Methodology, Investigation, S. G.; Writing, Review & editing, G.S.J.; Methodology, Formal Analysis, Writing, Review & Editing, Supervision, Funding Acquisition, J.B.; Conceptualization, Methodology, Writing, Review & Editing, Supervision, Funding Acquisition, A.B.

## Materials and methods

### Key resources table

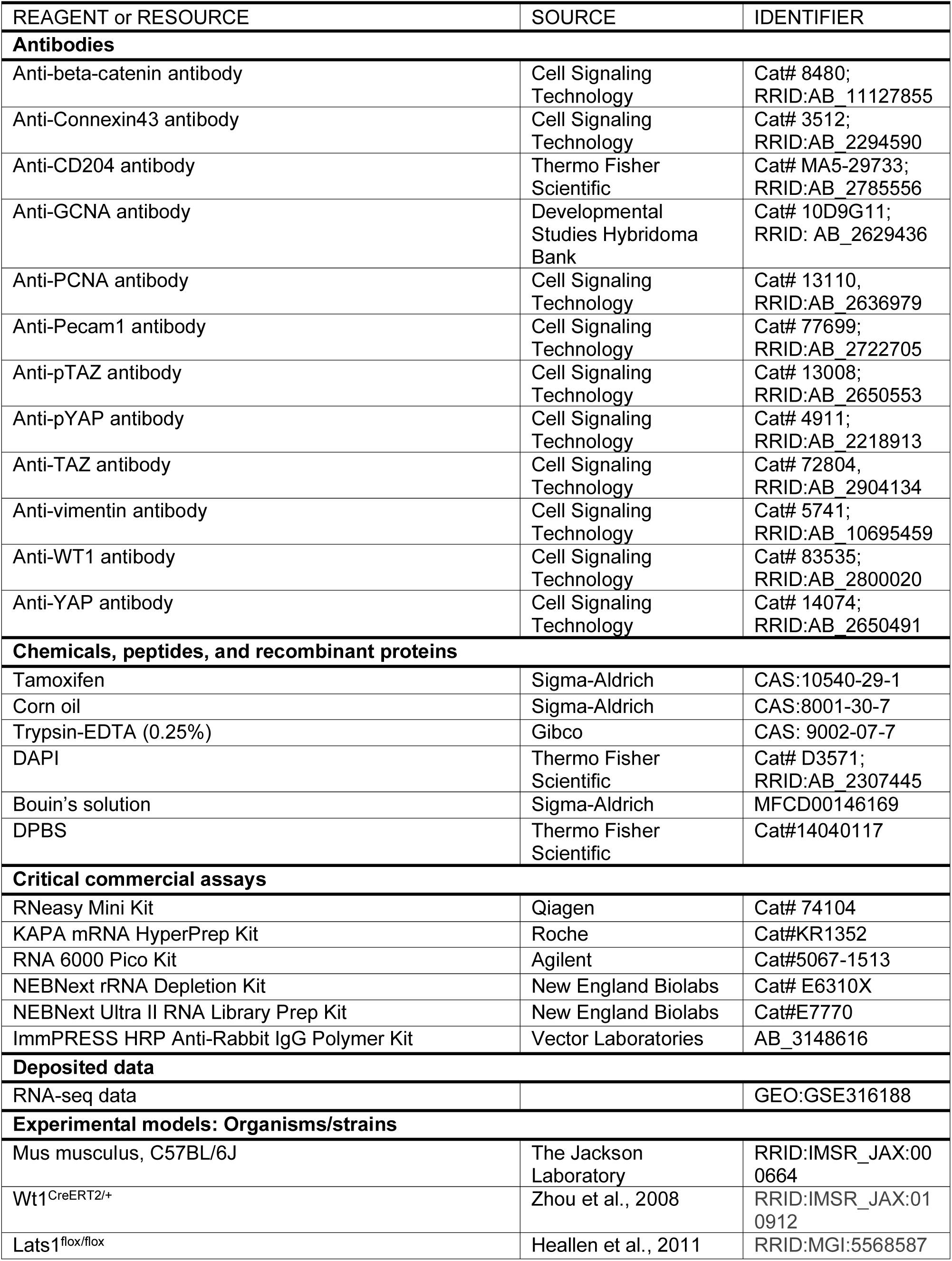
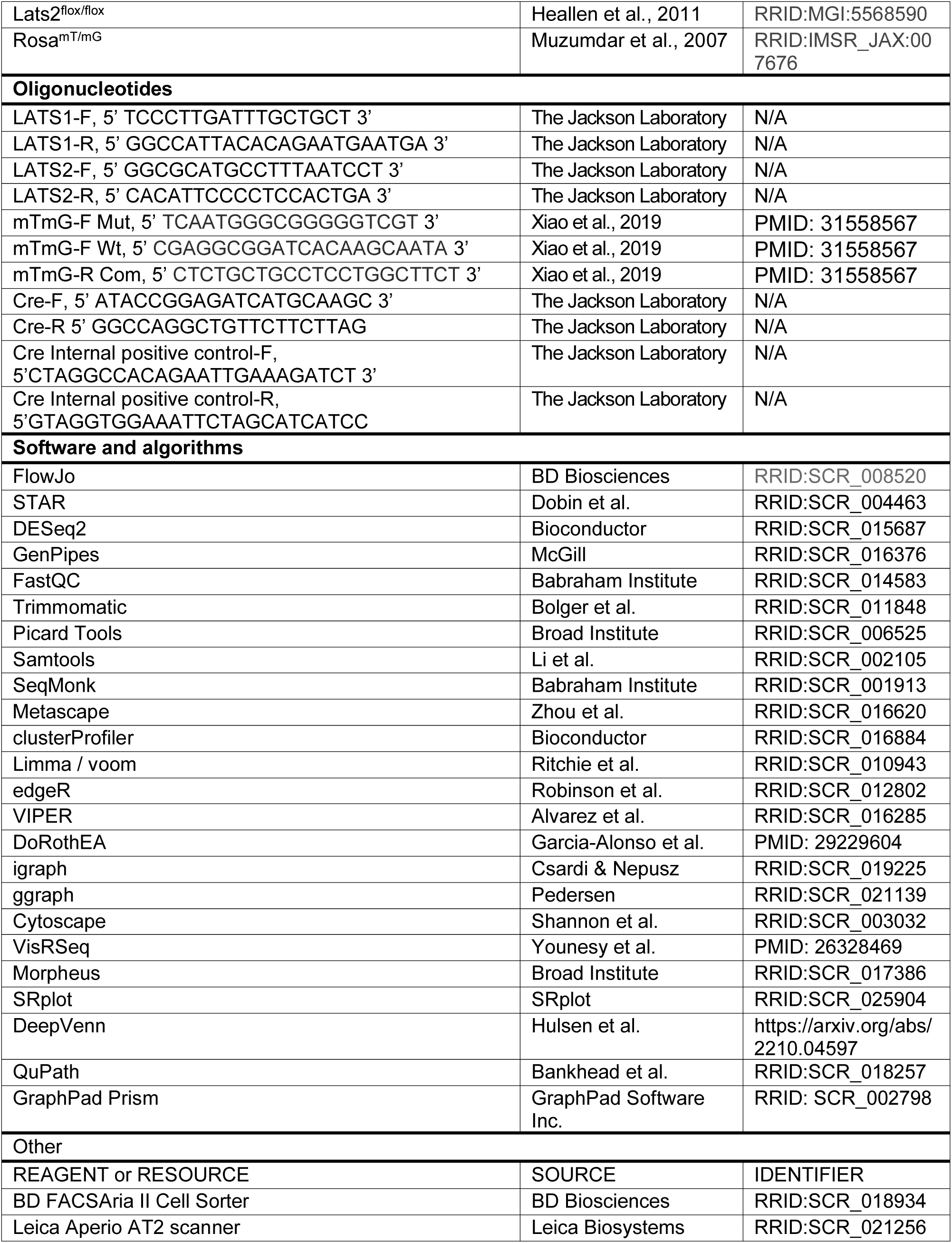

### Mice

All animal procedures were approved by the Comité d’Éthique de l’Utilisation des Animaux of the Université de Montréal (CÉUA; #Rech-1739/Rech-2081) and conducted in accordance with the guidelines of the Canadian Council on Animal Care (CCAC). Eight-week-old male C57BL/6J mice were used in all experiments. Animals were housed under standard conditions with ad libitum access to food and water. Group sizes and number of biological replicates are indicated in the corresponding figure legends.

## Mouse strains and genotyping

The following mouse lines were used : *Wt1*^CreERT2/+^ (Zhou et al., 2008), *Lats1*^flox/flox^ and *Lats2*^flox/flox^ (Heallen et al., 2013; Heallen et al., 2011) and *Rosa^mT/mG^* (Muzumdar et al., 2007), all maintained on a C57BL6/J background. Genotyping was performed by PCR on tail biopsies using primers and conditions previously described (Abou Nader et al., 2025; Ménard et al., 2020).

## Tamoxifen induction and experimental design

To induce Cre recombinase activity, eight-week-old *Lats1*^flox/flox^; *Lats2*^flox/flox^; *Wt1*^CreERT2/+^ mice and control littermates (*Lats1*^flox/flox^; *Lats2*^flox/flox^) received intraperitoneal injections of tamoxifen (120 mg/kg) or vehicle (corn oil) once daily for five consecutive days. Mice were sacrificed on days 1, 3, 5, and 12 following the final injection for time-course analyses. For fluorescence-activated cell sorting (FACS) experiments, Rosa*_mTmG/mTmG;_* Lats1_flox/flox;_ Lats2_flox/flox;_ Wt1_CreERT2/+_ and Rosa*_mTmG/mTmG;_* Wt1_CreERT2/+_ mice were used and sacrificed at day 12 post-induction. Vehicle-treated mice showed no observable differences compared with tamoxifen treated controls and were therefore not used for subsequent analyses.

## Tissue collection and histological processing

Mice were anesthetized by isoflurane inhalation and euthanized by cervical dislocation. Testes and epididymides were immediately dissected, weighed, and processed for histological analyses. Tissues were fixed overnight at 4°C in Bouin’s solution, dehydrated, and embedded in paraffin, and sectioned at 5 µm thickness. Sections were stained with hematoxylin and eosin (H&E) or Picrosirius Red or processed for immunohistochemistry.

## Histopathology, immunohistochemistry and cell quantification

Immunohistochemistry was performed using the ImmPRESS HRP horse anti-rabbit (Vector Laboratories) according to the manufacturer’s instructions. Primary antibodies used are listed in Table S1. Whole-slide images were acquired using an Aperio AT2 scanner (Leica Biosystems). Quantitative analyses were performed using QuPath v0.3.2. WT1+ cells were quantified across whole testis sections, whereas GCNA+ cells were quantified in a hundred seminiferous tubules. Two sections per mouse were analyzed. Data are presented as mean ± SEM from n = 6 mice per group.

## Fluorescence-activated cell sorting (FACS)

Rete testis was microdissected from the testes, rete testis and seminiferous tubules fragments were decapsulated and enzymatically digested with 0.25% trypsin-EDTA for 20 min at 37°C under agitation. Cell suspensions were filtered through a 70 µm strainer and centrifuged. Cells were resuspended in PBS supplemented with 2% fetal bovine serum and stained with DAPI (1 μg/mL). Cells were gated on singlets (FSC-A vs. FSC-H) and DAPI-negative to exclude dead cells. In Rosa^mTmG/mTmG^; Wt1^CreERT2/+^ animals, GFP-positive cells represented *Wt1*-lineage cells. Sorting was performed on a BD FACSAria II (BD Biosciences), and sorted cells were immediately processed for RNA extraction.

## RNA extraction and sequencing

### Whole Testis RNA Sequencing

Total RNA was extracted from whole testes using the RNeasy mini kit (Qiagen). Quality was assessed using an Agilent 2100 Bioanalyzer with the RNA 6000 Pico kit. Libraries (n = 3 control and n = 3 cKO littermates) were prepared using the KAPA mRNA Hyperprep kit (Roche). Single-read sequencing (75-85 bp) was performed on an Illumina NextSeq 500 platform, yielding 27-42 million reads per sample, Illumina) at the Genomics Core Facility of the Institute for Research in Immunology and Cancer.

## FACS-sorted cell RNA Sequencing

Total RNA was isolated from FACS-sorted cells obtained from control (*Rosa^mTmG/mTmG^; Wt1*^CreERT2/+^) and cKO (*Rosa^mTmG/mTmG^; Lats1*^flox/flox^; *Lats2*^flox/flox^; *Wt1*^CreERT2/+^) mice. Ribosomal RNA was depleted using the NEBNext rRNA Depletion Kit (#E6310X). Double-stranded cDNA synthesis and library preparation were performed using NEBNext Ultra II kits. Libraries were sequenced as 150 bp paired end on the NovaSeq 6000 platform at The Center for Applied Genomics (Toronto, Ontario, Canada).

## RNA-Seq data processing and analysis

### Whole testis RNA-Seq

Reads were aligned to the mouse genome (GRCm38, Gencode M23) using STAR (v 2.7.1a). Gene expression was quantified using SeqMonk v1.48.1. Differential expression analysis was performed using DESeq2 (v1.22.2) within SeqMonk with a significance threshold of p-value < 0.05 and fold change cutoff of ±1. Gene ontology enrichment analyses were conducted using Metascape. Data visualization was performed using VisRSeq, DeepVenn, Morpheus and SRplot. Germ cell-specific genes were identified based on a previously published gene list (Jung et al., 2019). Wilcoxon Signed Rank test (two-tailed, α = 0.05) was used to compare theoretical (0) and observed medians across eight spermatogenic stages. All comparisons were significant (p < 0.0001). Data are shown as median ± 95% confidence interval.

## FACS-sorted cell RNA-Seq

FASTQ files were processed using GenPipes v3.1.2. Quality control was performed with FastQC and reads were trimmed using Trimmomatic. Alignment to the mouse genome (mm10) was performed using STAR, followed by processing with Picard and Samtools. Differential expression analysis was conducted using DESeq2 within SeqMonk (p < 0.05, |log2FC| > 1). Gene ontology enrichment was performed using Metascape. Data visualization was performed using FlowJo, VisRSeq, DeepVenn, Morpheus, and SRplot. TF activity was inferred using VIPER with DoRothEA (A–C regulons), following TMM normalization and voom transformation. Differential TF activity was assessed using limma linear models with Benjamini–Hochberg correction (FDR < 0.05), including genotype-by-cell type interactions. Regulatory networks were constructed from significantly altered TFs and target genes showing concordant expression changes. TF–TF similarity was quantified using Jaccard overlap of regulons, filtered at the 90th percentile, and clustered using Louvain modularity. Functional annotation was performed using GO enrichment (clusterProfiler), with top terms defining module identity. Networks were visualized in R (igraph, ggraph) and Cytoscape.

## Quantification and statistical analysis

Statistical analyses were performed using Prism v9 (GraphPad). A two-tailed Student’s t-test were used for comparisons between two groups. Statistical significance was defined as p < 0.05. Data are presented as mean ± SEM.

---

**Supplemental Figure S1.**
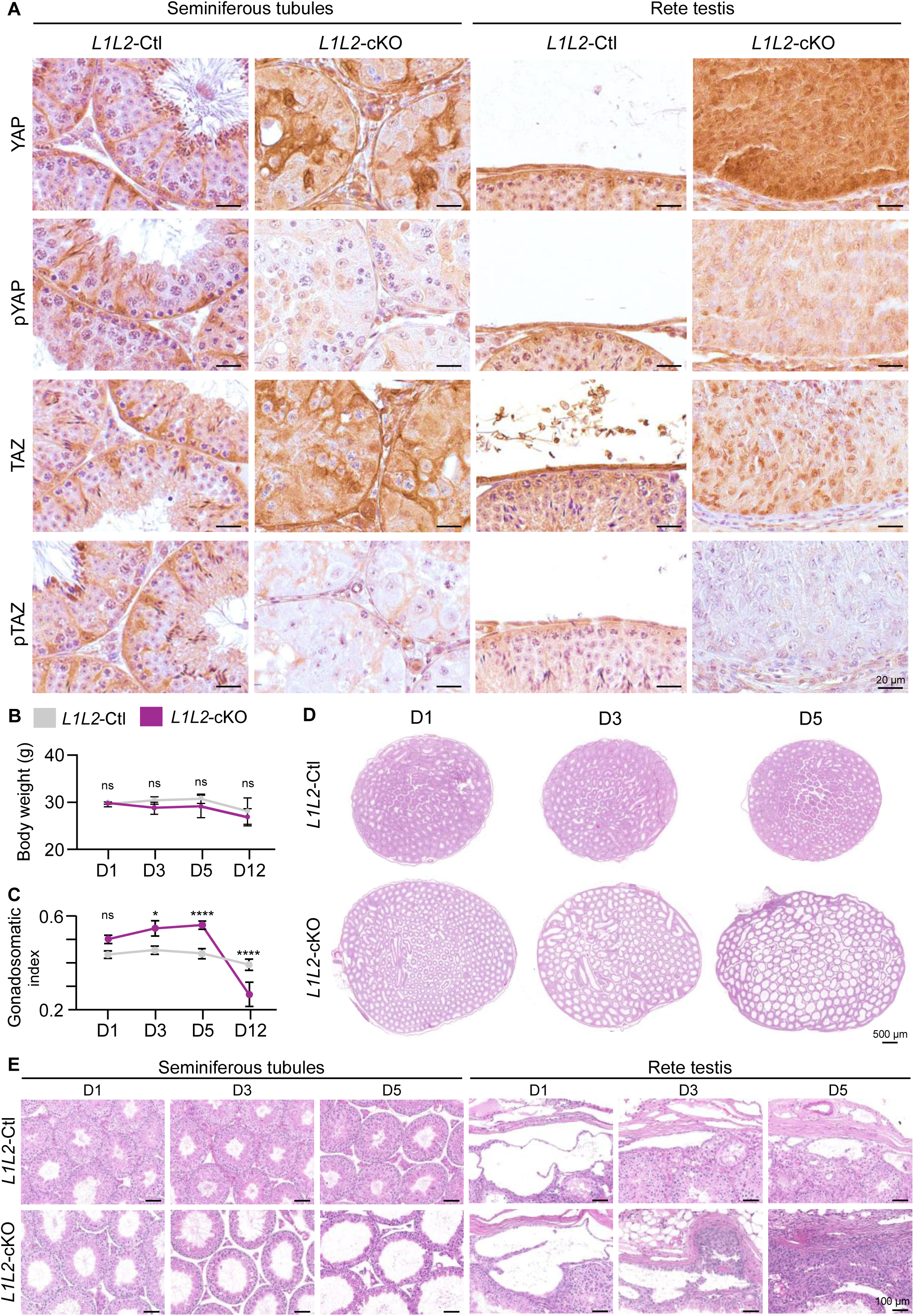
Validation of tamoxifen-inducible Lats1/2 deletion and Hippo pathway inactivation, related to. **Figure 1. (A)** IHC staining for YAP, TAZ and their phosphorylated forms (pYAP/pTAZ) in Sertoli and rete testis sections of *L1L2*-Ctl and *L1L2*-cKO testes at D12. Scale bars, 20 µm. **(B)** Time-course of body weight in *L1L2*-Ctl and *L1L2*-cKO mice between D1 and D12 (n = 6 per group). ns: not statistically significant, Student’s t-test. **(C)** Time-course comparing the gonadosomatic index in *L1L2*-cKO mice compared to *L1L2*-Ctl mice between D1 and D12 (n = 6 per group). *: p < 0.05, ****: p < 0.0001, ns: not statistically significant, Student’s t-test. **(D)** H&E staining in whole-testis cross-sections of *L1L2*-Ctl and *L1L2*-cKO testes between D1 and D12. Scale bars, 500 µm. **(E)** H&E staining in seminiferous tubules or rete testis sections from *L1L2*-Ctl and *L1L2*-cKO mice between D1 and D12. Scale bars, 100 µm.

**Supplemental Figure S2.**
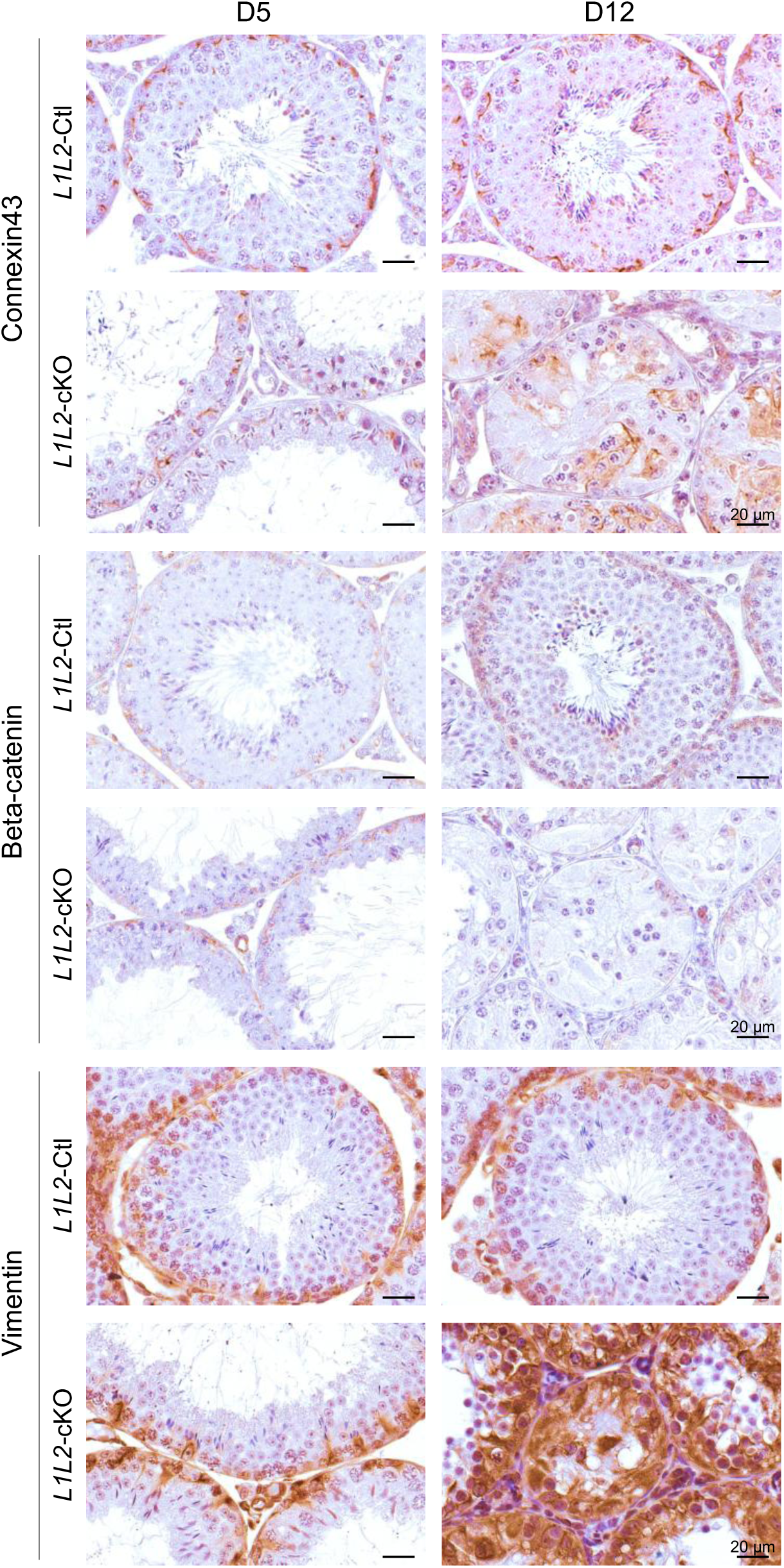
Altered junctional and cytoskeletal markers following *Lats1/2* deletion, related to. **Figure 2**. IHC for Connexin43, beta-catenin and vimentin in L1L2-Ctl and L1L2-cKO testes at D5 and D12. Scale bars, 20µm.

**Supplemental Figure S3.**
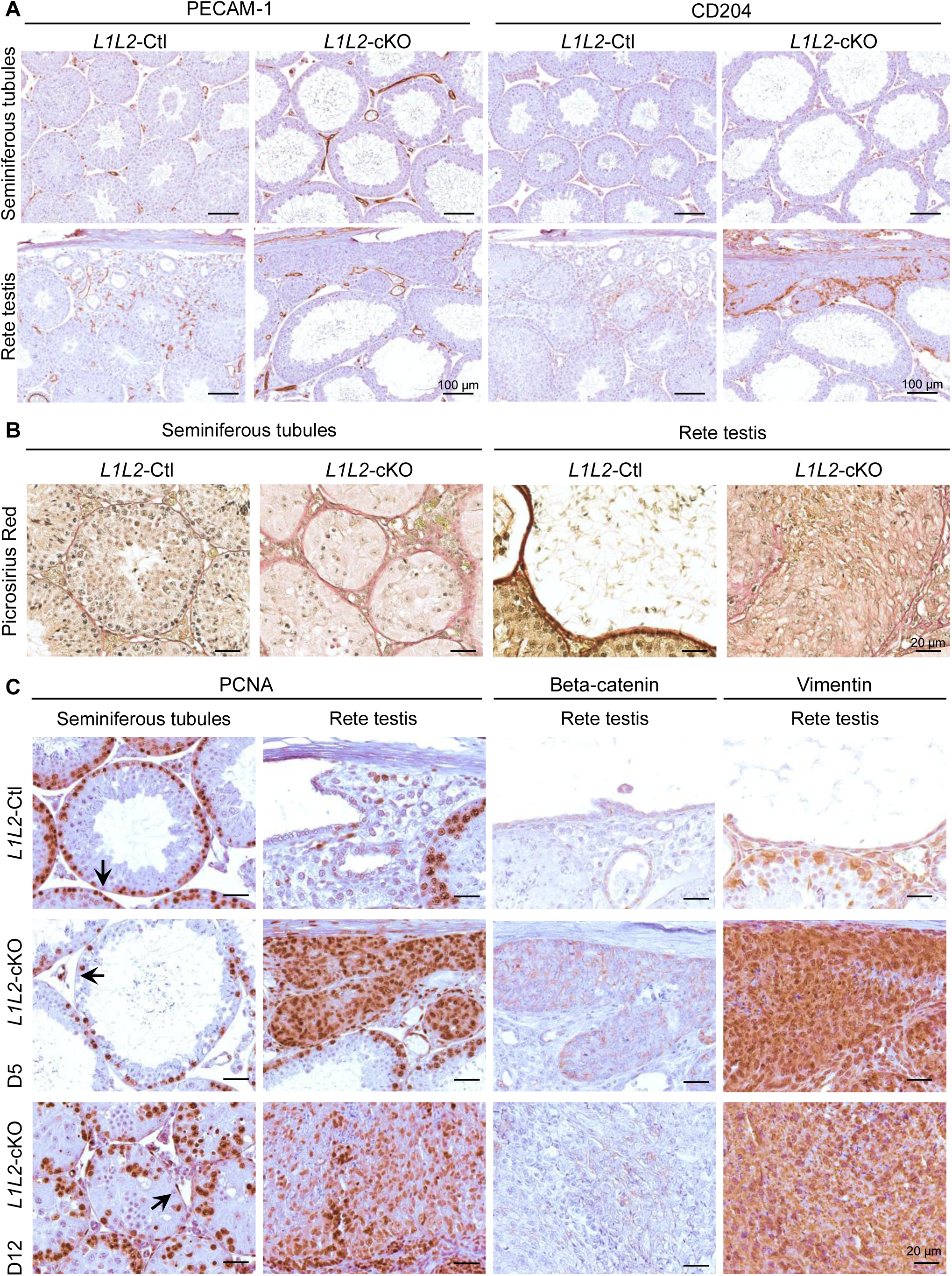
Early vascular and immune remodeling and rete testis proliferation, related to. **Figure 3. (A)** IHC for PECAM-1 and CD204 in seminiferous tubules and rete testis sections of *L1L2*-Ctl and *L1L2*-cKO testes at D5 and D12. Scale bars, 100µm. **(B)** Picrosirius Red in seminiferous tubules and rete testis sections of *L1L2*-Ctl and *L1L2*-cKO testes at D12. Scale bars, 20µm. **(C)** IHC for PCNA, beta-catenin and vimentin in seminiferous tubules and rete testis sections of *L1L2*-Ctl and *L1L2*-cKO mice at D5 and D12. Scale bars, 20µm. black arrow: Sertoli cells.

**Supplemental Figure S4.**
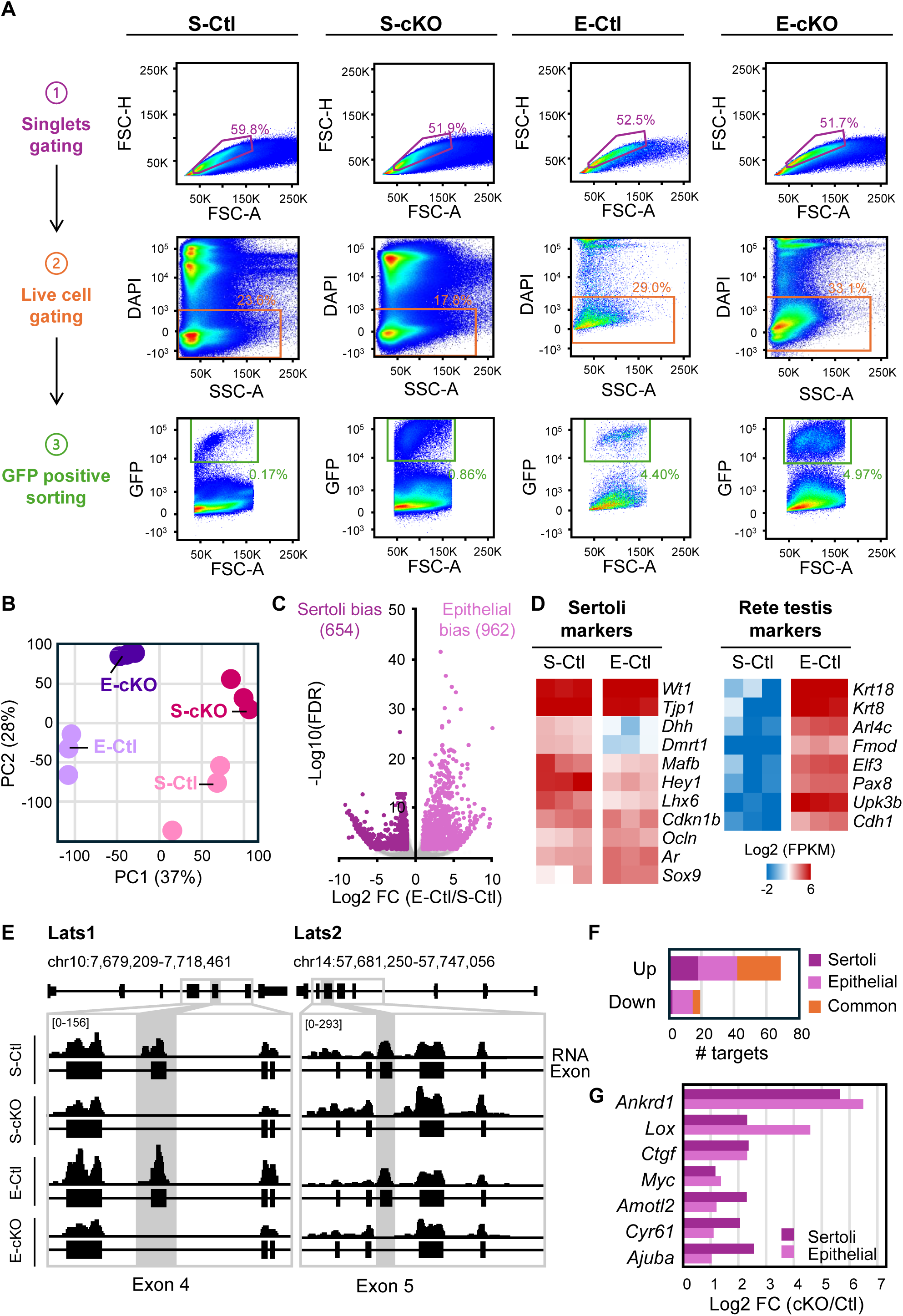
Flow-cytometric isolation and transcriptional validation of Sertoli and rete testis epithelial cells, related to. **Figure 4. (A)** Flow cytometry plots showing the gating strategy for eGFP+ sorting of WT1-lineage cells Sertoli (S) or rete testis epithelial (E) cells from *L1L2*-Ctl or *L1L2*-cKO testes at D12. **(B)** Principal component analysis (PCA) plot showing segregation of RNA-seq libraries by cell type and genotype (PC1 and PC2). **(C)** Volcano plot showing differentially expressed genes between Sertoli and rete testis epithelial WT1+ cells from *L1L2*-Ctl testes. **(D)** Expression of lineage-specific transcriptional signatures associated with Sertoli or rete testis epithelial cells. **(E)** Genome browser tracks showing RNA-seq coverage across *Lats1* and *Lats2* in Sertoli (S-Ctl, S-cKO) and rete testis epithelial (E-Ctl, E-cKO) cells. Exons targeted by the conditional knockout are highlighted in gray. **(F)** Stacked bar chart showing RNA-seq expression of selected YAP/TAZ target genes in Sertoli (S-Ctl, S-cKO) and rete testis epithelial (E-Ctl, E-cKO) cells. Colors indicate cell type specificity: Sertoli (purple), epithelial (pink), or shared (orange). **(G)** Bar chart showing differential expression of selected canonical YAP/TAZ target genes between S-cKO and S-Ctl or E-cKO and E-Ctl cells.

**Supplemental Figure S5.**
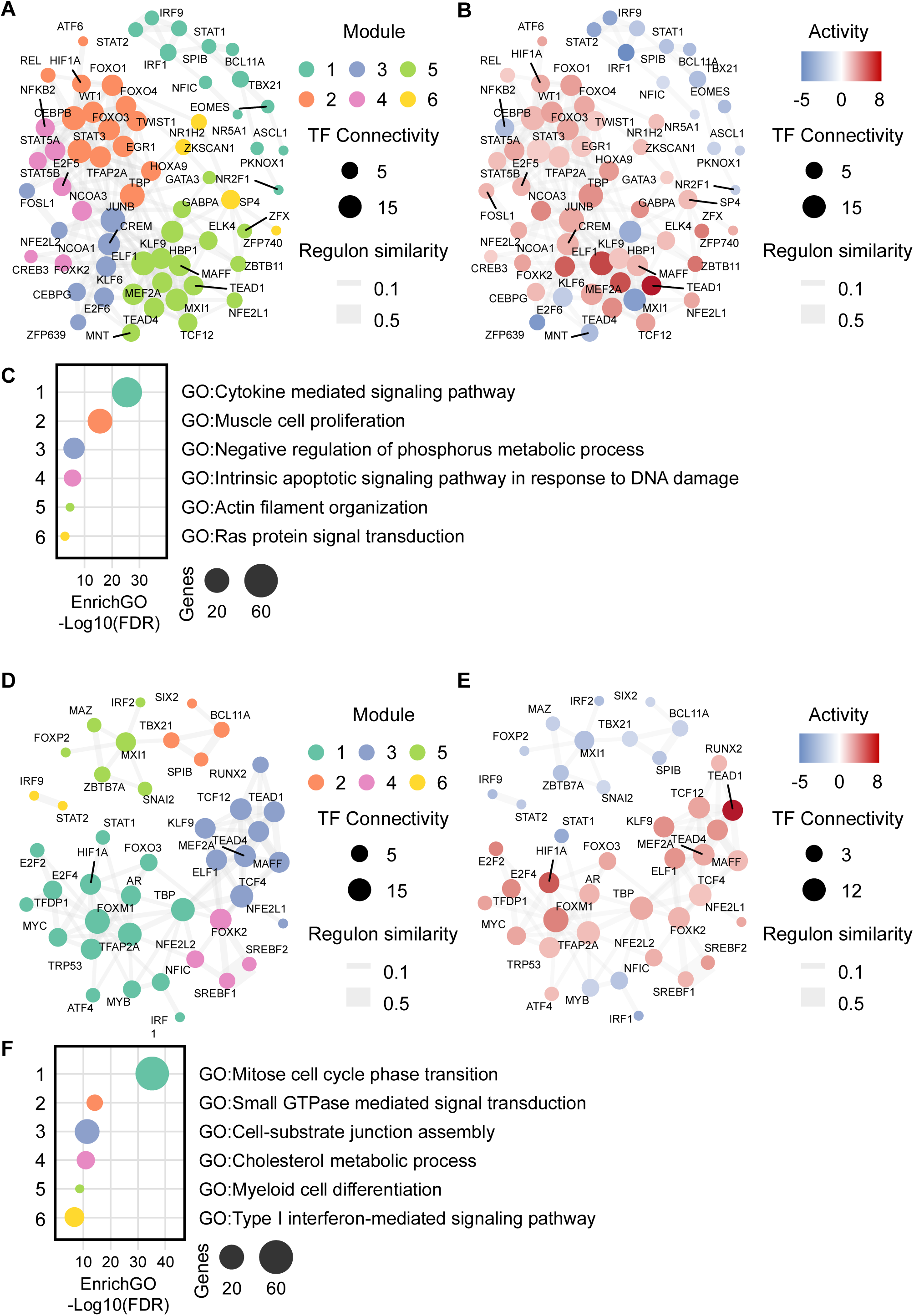
Regulatory network organization in Sertoli and rete testis epithelial cells following *Lats1/2* deletion, related to Figure 5. (A, D) Transcription factor regulatory networks reconstructed from VIPER-derived regulon similarity in Sertoli **(A)** and rete testis epithelial **(D)** cells. Edges represent regulon overlap (Jaccard index), node size reflects TF connectivity, and colors indicates Louvain-defined modules. **(B, E)** Reconstructed Transcription factor regulatory networks layout representing VIPER inferred TF activity in Sertoli **(B)** and rete testis epithelial **(E)** cells. **(C, F)** Gene Ontology enrichment analysis of modules identified in Sertoli **(C)** and rete testis epithelial **(F)** cells. Functional annotations were derived from the target genes associated with TF within each module.

## References

1. Abou Nader, N., Charrier, L., Meisnsohn, M. C., Banville, L., Deffrennes, B., St-Jean, G., Boerboom, D., Zamberlam, G., Brind’Amour, J., Pépin, D., & Boyer, A. (2024). Lats1 and Lats2 regulate YAP and TAZ activity to control the development of mouse Sertoli cells. Faseb j, 38(9), e23633. 10.1096/fj.202400346R

2. Abou Nader, N., Jakuc, N., Meinsohn, M. C., Charrier, L., Banville, L., Brind’Amour, J., Paquet, M., St-Jean, G., Boerboom, D., Mao, J., Pépin, D., Breault, D. T., Zamberlam, G., & Boyer, A. (2025). Hippo Signaling Is Essential for the Maintenance of Zona Glomerulosa Cell Fate in the Murine Adrenal Cortex. Endocrinology, 166(6). 10.1210/endocr/bqaf077

3. Ahmed, E. A. (2026). Sertoli cell aging: damage accumulation and epigenetic alterations affecting male fertility. Biogerontology, 27(2), 47. 10.1007/s10522-026-10392-6

4. Basten, S. G., & Giles, R. H. (2013). Functional aspects of primary cilia in signaling, cell cycle and tumorigenesis. Cilia, 2(1), 6. 10.1186/2046-2530-2-6

5. Carter, P., Schnell, U., Chaney, C., Tong, B., Pan, X., Ye, J., Mernaugh, G., Cotton, J. L., Margulis, V., Mao, J., Zent, R., Evers, B. M., Kapur, P., & Carroll, T. J. (2021). Deletion of Lats1/2 in adult kidney epithelia leads to renal cell carcinoma. J Clin Invest, 131(11). 10.1172/jci144108

6. Clulow, J., Jones, R. C., Hansen, L. A., & Man, S. Y. (1998). Fluid and electrolyte reabsorption in the ductuli efferentes testis. J Reprod Fertil Suppl, 53, 1–14. https://www.ncbi.nlm.nih.gov/pubmed/10645261

7. Cordenonsi, M., Zanconato, F., Azzolin, L., Forcato, M., Rosato, A., Frasson, C., Inui, M., Montagner, M., Parenti, A. R., Poletti, A., Daidone, M. G., Dupont, S., Basso, G., Bicciato, S., & Piccolo, S. (2011). The Hippo transducer TAZ confers cancer stem cell-related traits on breast cancer cells. Cell, 147(4), 759–772. 10.1016/j.cell.2011.09.048

8. Engeland, K. (2022). Cell cycle regulation: p53-p21-RB signaling. Cell Death Differ, 29(5), 946–960. 10.1038/s41418-022-00988-z

9. Goetz, S. C., & Anderson, K. V. (2010). The primary cilium: a signalling centre during vertebrate development. Nat Rev Genet, 11(5), 331–344. 10.1038/nrg2774

10. Guo, P., Wan, S., & Guan, K. L. (2025). The Hippo pathway: Organ size control and beyond. Pharmacol Rev, 77(2), 100031. 10.1016/j.pharmr.2024.100031

11. Heallen, T., Morikawa, Y., Leach, J., Tao, G., Willerson, J. T., Johnson, R. L., & Martin, J. F. (2013). Hippo signaling impedes adult heart regeneration. Development, 140(23), 4683–4690. 10.1242/dev.102798

12. Heallen, T., Zhang, M., Wang, J., Bonilla-Claudio, M., Klysik, E., Johnson, R. L., & Martin, J. F. (2011). Hippo pathway inhibits Wnt signaling to restrain cardiomyocyte proliferation and heart size. Science, 332(6028), 458–461. 10.1126/science.1199010

13. Hedger, M. P. (2011). Immunophysiology and pathology of inflammation in the testis and epididymis. J Androl, 32(6), 625–640. 10.2164/jandrol.111.012989

14. Jung, M., Wells, D., Rusch, J., Ahmad, S., Marchini, J., Myers, S. R., & Conrad, D. F. (2019). Unified single-cell analysis of testis gene regulation and pathology in five mouse strains. Elife, 8. 10.7554/eLife.43966

15. Levasseur, A., Paquet, M., Boerboom, D., & Boyer, A. (2017). Yes-associated protein and WW-containing transcription regulator 1 regulate the expression of sex-determining genes in Sertoli cells, but their inactivation does not cause sex reversal. Biol Reprod, 97(1), 162–175. 10.1093/biolre/iox057

16. Lopez-Hernandez, A., Sberna, S., & Campaner, S. (2021). Emerging Principles in the Transcriptional Control by YAP and TAZ. Cancers (Basel), 13(16). 10.3390/cancers13164242

17. Major, A. T., Estermann, M. A., & Smith, C. A. (2021). Anatomy, Endocrine Regulation, and Embryonic Development of the Rete Testis. Endocrinology, 162(6). 10.1210/endocr/bqab046

18. Ménard, A., Abou Nader, N., Levasseur, A., St-Jean, G., Le Roy, M. L. G., Boerboom, D., Benoit-Biancamano, M. O., & Boyer, A. (2020). Targeted Disruption of Lats1 and Lats2 in Mice Impairs Adrenal Cortex Development and Alters Adrenocortical Cell Fate. Endocrinology, 161(5). 10.1210/endocr/bqaa052

19. Misra, J. R., & Irvine, K. D. (2018). The Hippo Signaling Network and Its Biological Functions. Annu Rev Genet, 52, 65–87. 10.1146/annurev-genet-120417-031621

20. Mooring, M., Fowl, B. H., Lum, S. Z. C., Liu, Y., Yao, K., Softic, S., Kirchner, R., Bernstein, A., Singhi, A. D., Jay, D. G., Kahn, C. R., Camargo, F. D., & Yimlamai, D. (2020). Hepatocyte Stress Increases Expression of Yes-Associated Protein and Transcriptional Coactivator With PDZ-Binding Motif in Hepatocytes to Promote Parenchymal Inflammation and Fibrosis. Hepatology, 71(5), 1813–1830. 10.1002/hep.30928

21. Moya, I. M., & Halder, G. (2019). Hippo-YAP/TAZ signalling in organ regeneration and regenerative medicine. Nat Rev Mol Cell Biol, 20(4), 211–226. 10.1038/s41580-018-0086-y

22. Muzumdar, M. D., Tasic, B., Miyamichi, K., Li, L., & Luo, L. (2007). A global double-fluorescent Cre reporter mouse. Genesis, 45(9), 593–605. 10.1002/dvg.20335

23. O’Donnell, L., Smith, L. B., & Rebourcet, D. (2022). Sertoli cells as key drivers of testis function. Semin Cell Dev Biol, 121, 2–9. 10.1016/j.semcdb.2021.06.016

24. Panciera, T., Azzolin, L., Cordenonsi, M., & Piccolo, S. (2017). Mechanobiology of YAP and TAZ in physiology and disease. Nat Rev Mol Cell Biol, 18(12), 758–770. 10.1038/nrm.2017.87

25. Sen Sharma, S., Vats, A., & Majumdar, S. (2019). Regulation of Hippo pathway components by FSH in testis. Reprod Biol, 19(1), 61–66. 10.1016/j.repbio.2019.01.003

26. Siu, M. K., & Cheng, C. Y. (2008). Extracellular matrix and its role in spermatogenesis. Adv Exp Med Biol, 636, 74–91. 10.1007/978-0-387-09597-4_5

27. Theas, M. S. (2018). Germ cell apoptosis and survival in testicular inflammation. Andrologia, 50(11), e13083. 10.1111/and.13083

28. Totaro, A., Panciera, T., & Piccolo, S. (2018). YAP/TAZ upstream signals and downstream responses. Nat Cell Biol, 20(8), 888–899. 10.1038/s41556-018-0142-z

29. Turinsky, L., Rehrl, S., Nguyen, C., Benyahia, S., Kuperwasser, N., Vasseur, F., Arevalo, A. R., Rabant, M., Goudin, N., Mesnard, L., Canaud, G., Pontoglio, M., Isnard, P., & Terzi, F. (2025). Podocyte-specific YAP activation triggers crescentic glomerulonephritis through cross-talk with parietal epithelial cells. Sci Transl Med, 17(804), eadq7825. 10.1126/scitranslmed.adq7825

30. Walker, W. H. (2003). Molecular mechanisms controlling Sertoli cell proliferation and differentiation. Endocrinology, 144(9), 3719–3721. 10.1210/en.2003-0765

31. Weigel Munoz, M., Cohen, D. J., Da Ros, V. G., Gonzalez, S. N., Rebagliati Cid, A., Sulzyk, V., & Cuasnicu, P. S. (2024). Physiological and pathological aspects of epididymal sperm maturation. Mol Aspects Med, 100, 101321. 10.1016/j.mam.2024.101321

32. Wheway, G., Nazlamova, L., & Hancock, J. T. (2018). Signaling through the Primary Cilium. Front Cell Dev Biol, 6, 8. 10.3389/fcell.2018.00008

33. Wicks, E. E., & Semenza, G. L. (2022). Hypoxia-inducible factors: cancer progression and clinical translation. J Clin Invest, 132(11). 10.1172/JCI159839

34. Zanconato, F., Forcato, M., Battilana, G., Azzolin, L., Quaranta, E., Bodega, B., Rosato, A., Bicciato, S., Cordenonsi, M., & Piccolo, S. (2015). Genome-wide association between YAP/TAZ/TEAD and AP-1 at enhancers drives oncogenic growth. Nat Cell Biol, 17(9), 1218–1227. 10.1038/ncb3216

35. Zhao, S., Zhu, W., Xue, S., & Han, D. (2014). Testicular defense systems: immune privilege and innate immunity. Cell Mol Immunol, 11(5), 428–437. 10.1038/cmi.2014.38

36. Zhou, B., Ma, Q., Rajagopal, S., Wu, S. M., Domian, I., Rivera-Feliciano, J., Jiang, D., von Gise, A., Ikeda, S., Chien, K. R., & Pu, W. T. (2008). Epicardial progenitors contribute to the cardiomyocyte lineage in the developing heart. Nature, 454(7200), 109–113. 10.1038/nature07060

37. Zhu, Y., Abedini, A., Rodriguez, G. M., McCloskey, C. W., Abou-Hamad, J., Salah, O. S., Larocque, J., Tsoi, M. F., Boerboom, D., Cook, D., & Vanderhyden, B. (2025). Loss of LATS1 and LATS2 promotes ovarian tumor formation by enhancing AKT activity and PD-L1 expression. Oncogene, 44(27), 2240–2252. 10.1038/s41388-025-03387-z

